# Wavelet-based Approach for Diagnosing Attention Deficit Hyperactivity Disorder (ADHD)

**DOI:** 10.1101/2022.10.04.510864

**Authors:** Dixon Vimalajeewa, Ethan McDonald, Scott Alan Bruce, Brani Vidakovic

**Affiliations:** Texas A&M University

## Abstract

Attention deficit hyperactivity disorder (ADHD) is a common cognitive disorder affecting children. ADHD can interfere with educational, social, and emotional development, so early detection is essential for obtaining proper care. Standard ADHD diagnostic protocols rely heavily on subjective assessments of perceived behavior. An objective diagnostic measure would be a welcome development and potentially aid in accurately and efficiently diagnosing ADHD. Analysis of pupillary dynamics has been proposed as a promising alternative method of detecting affected individuals effectively. This study proposes a method based on the self-similarity of pupillary dynamics and assesses its strength as a potential diagnostic biomarker. Localized discriminatory features are developed in the wavelet domain and selected via a rolling window method to build classifiers. The application on a task-based pupil diameter time series dataset of children aged 10-12 years shows that the proposed method achieves greater than 78% accuracy in detecting ADHD. Comparing with a recent approach that constructs features in the original data domain, the proposed wavelet-based classifier achieves more accurate ADHD classification with fewer features. The findings suggest that the proposed diagnostic procedure involving interpretable wavelet-based self-similarity features of pupil diameter data can potentially aid in improving the efficacy of ADHD diagnosis.

## 1 Introduction

Attention deficit hyperactivity disorder (ADHD) is a common neurodevelopmental disorder among children characterized by a combination of symptoms, including decreases in attention and increases in hyperactivity and impulsivity. According to the Centers for Disease Control and Prevention (CDC), the majority of children diagnosed with ADHD are ages 6-12 years, but children aged 13-17 years, and even older young adults are also diagnosed with the disorder. Worldwide, the prevalence of ADHD among children and teenagers is approximately 5-8%, and more than 1 million children per year are diagnosed with ADHD in the United States alone [2]. This disorder can have a significant negative impact on educational, social, and emotional development which can lead to anxiety, depression, and antisocial behaviors [30]. Thus, early detection of ADHD for ensuring proper treatment is critical to promote typical development and reduce the risk of developing other psychological disorders.

Since there are a broad range of symptoms that can be associated with ADHD, the majority of current early diagnostic techniques rely largely on qualitative assessments of perceived behavior. For example, in the study by Gathje et al. [7], ADHD diagnoses were conducted using a checklist of 18 symptoms related to hyperactivity and impulsivity. Assessments required respondents to characterize the frequency and severity of particular behaviors using subjective ordinal scales (e.g. four point scale: “not at all”, “just a little”, “pretty much” or “very much”; nine point scale from “mild” to “severe”). Subjective behavioral assessments are prone to inconsistency associated with different perceptions of frequency and severity, bias due to greater familiarity with recent behavior, and potentially incomplete recollection of past behaviors. These limitations may result in significant risk of over- or under-diagnosis of ADHD [5].

In order to provide a more standardized and objective characterization of ADHD-related behaviors, quantitative methods have been proposed. These methods are based on physiological biomarkers, such as regional cerebral blood flow, pupil diameter, EEG (electroencephalogram), and blinking rate. For example, pupillary dynamics (i.e. high frequency changes in pupil diameter) have been proposed as a promising biomarker for accurate and efficient detection of ADHD. Pupillary dynamics provides real-time information about mental workload without disrupting the user’s ongoing activity (e.g., visual stimuli). Also, measuring pupillary dynamics is less intrusive compared to measuring other biomarkers. For example, Sou et al. [19] show that large pupil diameters, low temporal complexity, and symmetry are the three main properties associated with the pupil diameter in individuals with ADHD. Meanwhile, the study in Wainstein et al. [30] demonstrates that the pupil size can be used as a potential biomarker to detect ADHD. Besides pupil dynamics, EEG signals have also been used for diagnosing ADHD. A visualization method proposed in Krishna [14] utilizes EEG signals to identify abnormalities in different regions of the brain where there is excessive alpha wave activity to detect ADHD in children and adults. This study demonstrates that excessive frontal alpha activity in the brain is often associated with ADHD and depression.

These studies generally evaluate specific patterns after a series of data pre-processing steps (e.g. normalization, denoising) to generate potentially discriminatory biomarkers. Then, various feature selection and classification methods are applied to the features to build a model for detecting ADHD. Since there are variety of ways to perform these pre-processing steps, the features identified as predictive of ADHD could vary depending on the order and nature of the pre-processing steps taken, which can hamper reproducibility. Thus, approaches that rely on minimal data pre-processing would be desirable in order to simplify implementation and improve generalizability of findings. In addition, the studies reported in the literature [30, 20, 5] do not consider variability in associations among pupil diameter values that could indicate the presence of ADHD. As reflected in the literature [29], assessing self-similarity in pupil diameter data could effectively characterize such differences in associations with minimal pre-processing.

Self-similarity in a signal or time series is the phenomenon of exhibiting similar properties and behaviors when inspected at different resolutions. As reflected by vibrant research, self-similar properties are used frequently in disease diagnostics, and wavelets are suitable for modeling self-similarity. For example, to detect plasma cysteine deficiency, the self-similar properties of Nuclear Magnetic Resonance spectra are examined using wavelets in Jung et al. [11] Similarly, the study in Vimalajeewa et al. [29] reports the significance of using wavelet-based features to characterize self-similar behavior in protein mass spectra for ovarian cancer diagnosis. ADHD diagnosis studies in the published literature have also used wavelets, but mainly as a tool for pre-processing. For example, wavelets are used to denoise EEG signals for detecting ADHD-affected children in Lee et al [15]. Similarly, a wavelet-based method is used in Shi et al. [21] to improve classification performance of multiscale Schur monotone measures employed to analyze pupillary behaviors of individuals with ocular disease. However, to the best of our knowledge, no studies have explored self-similarity in pupillary dynamics. Thus, the novelty of this study is the use of self-similar behavior in pupil dynamics in the context of diagnosing ADHD.

The main contribution of this study is to propose a method to explore self-similar properties in pupillary dynamics and assess its strength as a potential discriminatory biomarker for ADHD diagnosis. Quantitative self-similarity analysis generally includes assessing scaling behavior (i.e. regularity) in signals at different scales (i.e. resolutions). This study explores self-similarity in pupil diameter data in the wavelet domain. More specifically, we compute wavelet spectra of the pupil diameter time series to characterize the regularity of level-wise decay in scale-specific energies of the resulting wavelet coefficients from the wavelet decomposition. The rate of this energy decay (i.e. slope) quantifies the degree of regularity in pupil diameter time series and is used as a discriminatory descriptor. The energies at different wavelet decomposition levels are computed as the variance of the wavelet coefficients. However, the traditional variance is sensitive to noise, and outliers frequently seen in pupil diameter data can negatively impact diagnostic performance; pupillary dynamics are generally observed as seemingly erratic, noisy, and high-frequency time series. To address this issue, the present study uses distance variance[3] to estimate level-wise wavelet energies because it is robust to noise and outliers and has not been previously used for characterizing wavelet spectra. Instead of relying on a single slope to characterize the entire pupil diameter time series, we use a rolling window-based approach. This rolling window approach enables exploring evolutionary slopes and extracting localized features that can be useful for subsequent detection of ADHD. The strength of those features in detecting ADHD is assessed with three popular classification algorithms: logistic regression (LR), support vector machine (SVM), and k-nearest neighbor (KNN). Commonly used key medical diagnostic classification performance metrics, sensitivity, specificity, and correct classification accuracy, are reported to assess the performance. We illustrate and evaluate ADHD detection performance using self-similarity features using a pupil-diameter dataset[20]. Our evaluation demonstrates improved diagnostic performance compared to an existing technique in the literature, the feature engineering-based classification method which is applied on the same dataset.

### Dataset

This study uses a publicly available dataset, featured in Rojas-Líbano et al. [20], consisting of pupil diameter time series collected from 50 children aged 10-12 (28 ADHD-diagnosed and 22 controls) during a visuospatial working memory task. A subgroup of 17 ADHD-diagnosed children performed the task twice, with and without administration of the medication methylphenidate. Thus, there are three pupil diameter data groups in the dataset: ADHD-diagnosed children with and without medication (*ADHD-on-medication* and *cases*) and children who are not diagnosed with ADHD.

In the visuospatial memory task, pupil diameter was recorded while a series of images was displayed on a screen in a systematic manner under controlled experimental settings. The *Eyelink 1000* device was used to record pupil diameter as well as time, location (x,y coordinates), and feedback at 1-kHz frequency. The task started by displaying a black cross at the center of the screen for 500*ms* (fixation window). Second, an image containing one or two dots located in a cell of the 4 *×* 4 grid was displayed on the screen for 750*ms*, followed by a fixation-time window. This process was repeated three times before displaying a distractor image for 1, 500*ms* in the third step. The distractor image was one of the four images, a task-related image (i.e., an image containing one or two dots located in any of the 4 *×* 4 grid), natural, emotional, none (i.e., no distractor image). In the fourth step, after a displaying a fixation-time window, a probe image was displayed for 1, 500*ms*. Finally, the participant was given 1, 500*ms* to give feedback (yes or no) whether the probe image was seen before the distractor image. This whole process is called a trial; on average, each participant took around 8 seconds to complete a trial. Each participant performed eight sessions, each containing 20 trials (i.e., 160 trials per participant). More details about the experiment and the dataset are available in Rojas-Líbano et al. [20].

Figure 1a, for example, shows a sample pupil-diameter dataset collected over one trial, 20 trials (one session), and an experiment (eight sessions). The pupil diameter data depicts highly spiky behavior, which may correspond to eye blinks, and contains many missing values. Also, the variability in pupil diameters at each time point (averaged over the total 160 trials under light intensity controlled environment) of the three groups (Fig. 1b) shows noticeable differences over time. Thus, this reflects the potential to differentiate ADHD-diagnosed children from the controls based on variability in pupil diameter.

**Figure 1:**
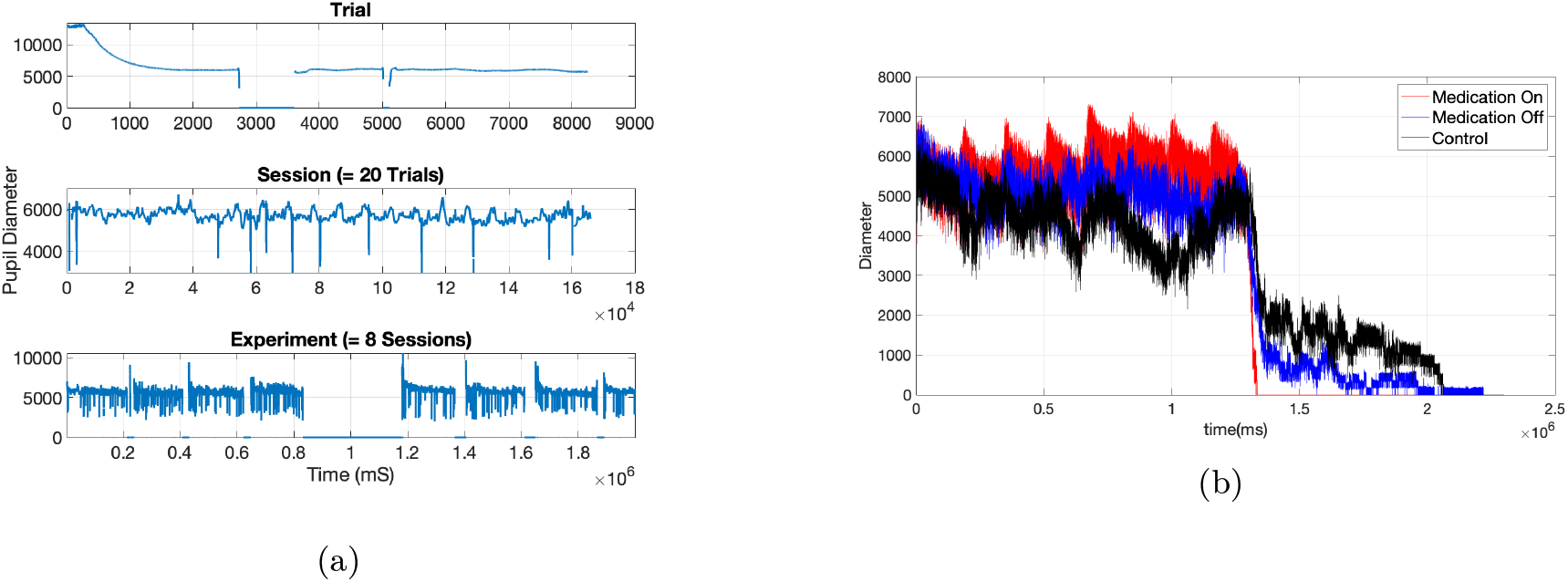
A sample pupil diameter dataset collected over a trial, session and an experiment (8 sessions) (a) and pupil diameter of the three groups (*ADHD-on-medication, ADHD-off-medication*, and *controls*) averaged over the number of 160 trials (b).

This dataset, however, has not yet been used extensively for ADHD diagnosis, and only a few studies have been reported in the literature. For example, in the study by Wainstein et al. [30] used basic statistical measures, such as minimum, maximum, correlation coefficient, and standard deviation, to characterize properties of the dataset. The data analysis outcomes of this study revealed that the *ADHD-off-medication* group indicated a decreased pupil diameter compared to the *ADHD-on-medication* group. The study by Das and Khanna [5] also reported an application of this dataset, proposing a feature engineering method-based machine-learning framework for automated detection of ADHD. This framework is the first machine learning-based approach reported in the literature using this dataset. These two studies, however, used only the pre-processed data (i.e. filtered, interpolated). Thus far, no study has considered the information hidden in the raw pupil diameter data or the information that might have been lost during data pre-processing. In the following, we present our wavelet-based self-similarity behavior analysis method and investigate its performance in effectively detecting ADHD.

## Methods

### Wavelet Transforms

Wavelet transforms (WTs) are standard tools in signal processing. The application of WTs on a signal transforms the signal into a set of contributions that are localized both in time and frequency. In the wavelet domain, the transformed signal has a hierarchical representation and allows for localized analysis at different scales (or resolutions) simultaneously. Thus, WTs enable assessment of certain inherent properties of signals, which are difficult to reveal in the original data domain. For example, scaling behavior of the signal energy is one of the properties that can be used to characterize different scale-sensitive behaviors such as long memory, fractality, and self-similarity[27].

The discrete wavelet transform (DWT), a popular choice of WT, has emerged as a tool of choice for analyzing complex signals in application domains where discrete data are analyzed. Simply, DWTs are linear transforms that can be represented as follows. Suppose a data vector *Y* = (*y*(*t*_1_), *y*(*t*_2_), *…, y*(*t*_*N*_))^*t*^ of size *N ×*1 is measured at *N* equally spaced points *t*_*i*_ for *i* = 1, 2, *…, N*. The DWT of *Y* is given by

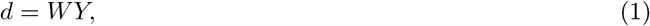

where *d* is also a vector of size *N ×* 1, and *W* is a wavelet-specific orthogonal matrix of size *N × N*. The elements in *W* are determined by the wavelet filter corresponding to the selected wavelet basis, such as the Haar, Daubechies, or Symmlet families[27].

Performing the DWT in matrix form, as in (1), is computationally expensive when *N* is large. A computationally fast algorithm developed by Mallat is used to perform the DWT for sample sizes that are a power of 2, e.g., *N* = 2^*J*^, *J ∈* Z^+^. The algorithm has a hierarchical structure, in which a series of convolutions are performed with a wavelet-specific low-pass filter and its counterpart, high-pass filter, followed by a decimation (i.e. selection of values at even positions in a sequence). This results in a multiresolution representation of the signal that comprised of a smooth approximation 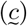 and a hierarchy of detail coefficients *d*_*jk*_ at different resolution levels (i.e. scale indexes) *j* and locations *k* within the same resolution level. Thus, for a vector *Y* of size *N* = 2^*J*^, the vector *d* in (1), has the following structure

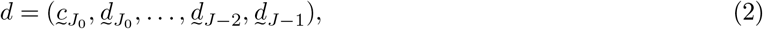

Where 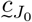 is a vector of coefficients corresponding to a smooth trend in the signal, 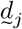 are detail coefficients at different resolutions *j*, and *J*_0_ is the coarsest resolution level in the wavelet decomposition, such that 1 ≤ *J*_0_ ≤ *J* − 1. Vector *d* in (2) has *J* − *J*_0_ levels of detail, 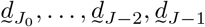, which are used in the definition of wavelet spectra. This algorithm is now available in many standard wavelet packages (e.g., *WAVELAB* module for MATLAB). Interested readers can find more details about WTs and different applications of WTs in Vidakovic [27].

### Self-similarity

Unlike any regular objects such as smooth curves and surfaces, many high-frequency signals acquired from sources in real-world processes, such as medicine, economics, and biology, are highly irregular. Self-similarity is a prominent property of such signals and simply defined as the nature of exhibiting similar properties and statistical behavior when inspected at different scales. That is, self-similar signals maintain either deterministic or statistical characteristics invariant to scaling. The Hurst exponent (*H*) is used to quantify self-similarity. It can also be understood as quantification of strength of dependence (or correlation structure) in a signal. A higher *H* value of a signal indicates that the signal exhibits a more regular and less erratic behavior. In general, *H <* 0.5 defines an anticorrelated signal, *H* = 0.5 is a random noise, and *H >* 0.5 indicates a positively correlated signal (or characterizes a long-memory process) [1]. Methods used to estimate *H* include Fourier-spectra methods, variance plots, quadratic variations, and zero-level crossings [12]. In particular, wavelet-based methods have enjoyed success in assessing and modeling self-similarity. In a more general setting, readers can find more details about self-similarity in Lee [16].

### Wavelet Spectra-based Self-Similarity Assessment

In assessing self-similarity, wavelet-based methods generally use wavelet spectra. A wavelet spectrum characterizes the relationship between the scale index *j* and energies at these scales. Energies are computed as the average magnitude of squared wavelet coefficients. Theoretically, wavelet coefficients have zero mean, so the average-square of wavelet coefficients is equivalent to the variance. Thus, the wavelet spectra can also be described as the logarithm for basis 2 of sample variance of wavelet coefficients in level *j*, as a function of that level,

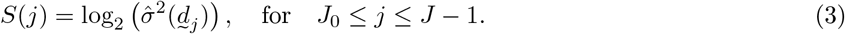

Here 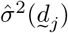 is an estimator of sample variance of vector 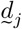. The spectral slope (and Hurst exponent) is estimated from the regression on pairs (*j, S*(*j*)).

Readers can find more details about the wavelet spectra and Hurst exponent, and their applications in Hamilton et al. [8] and Kong and Vidakovic [13].

The slope in a linear fit of pairs (*j, S*(*j*)), *J*_0_ ≤ *j* ≤ *J* − 1, is connected with a degree of regularity of the signal and used to calculate the Hurst exponent (*H*). Several types of regression of log energies *S* on the scale indices *j* are typically used for estimation of the slope. The Hurst exponent *H* is then computed as *H* = −(*slope* + 1)*/*2. Different techniques exist to estimate slope, and they may rely on some other measures, usually robust ones (Kang and Vidakovic [12]; Feng and Vidakovic [6]). This study, however, introduces a *distance variance*-based method for robustification of the slope estimator.

### Distance Variance-based Wavelet Spectra

The distance variance(covariance) is a robust estimator of the population variance(covariance) and has been widely applied to achieve robustness when data contain noise and outliers. For example, the studies Székely and Rizzo [23], Tsyawo and Soale [26], and Li et al. [17] have used the distance covariance to robustify the detection of non-linearity and selection of informative descriptors in feature screening, respectively. In Matteson and Tsay [18], the distance covariance was applied to perform independent component analysis (ICA) and achieved higher performance compared to the existing ICA methods such as, FastICA and kernel density-based ICA. The study in Cowley et al. [4] also demonstrates the use of distance covariance in the context of dimensionality reduction. The authors report that the distance covariance achieves better (or comparable) performance, compared to the two existing dimensionality reduction techniques, principal component analysis (PCA) and canonical component analysis (CCA). The distance variance, however, has not yet been applied in order to compute wavelet spectra. In this study, the levelwise variance of wavelet coefficients is estimated by the distance variance of wavelet coefficients to compute wavelet spectra. The distance variance of wavelet coefficients is computed as follows.

Suppose 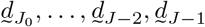 represents a vector of wavelet coefficients at a certain multiresolution level of signal *Y*. Let *a*_*ij*_ = |*d*_*i*_ − *d*_*j*_|, for *i, j* = 1, 2, *…, n* denotes the absolute difference between paired elements in 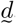. Then, given the double-centered difference *A*_*ij*_ as

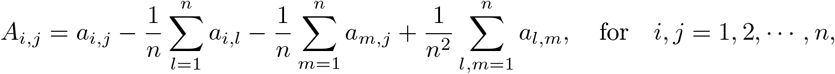

the distance variance of is defined as

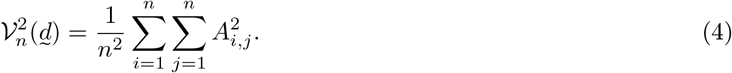

Readers can find more theoretical information about the distance variance(covariance) and applications in Székely and Rizzo [24], [22].

The distance variance is an outlier-resistant and statistically stable measure of variability. To illustrate this, we conducted an experiment on the impact of noise (or outliers) on the standard variance and distance variance in estimating the Hurst exponent of a Brownian motion using wavelet-based spectra. The standard Brownian motion was used to generate a data sample of size 1024. Theoretically, the Hurst exponent of this data is 0.5. A wavelet transform was applied to the data with Daubechies-6 wavelet and decomposition level 9. At each decomposition level, a set of 100 wavelet coefficients was randomly selected and contaminated by adding Gaussian noise. Then, wavelet spectra of the contaminated sample was computed using the standard variance and distance variance, and the Hurst exponents were found. The Hurst exponent estimated from the distance variance-based method deviates less from the theoretical Hurst exponent (*H* = 0.5), compared to the calculations using the standard sample variance (Fig. 2). Also, the spread of the estimated *H* values in the distance variance-based method is smaller. Thus, the estimation based on distance variance is less affected by contaminating a selfsimilar signal by the white noise. Similar stability in estimation can be obtained if outliers are added to the original data.

**Figure 2:**
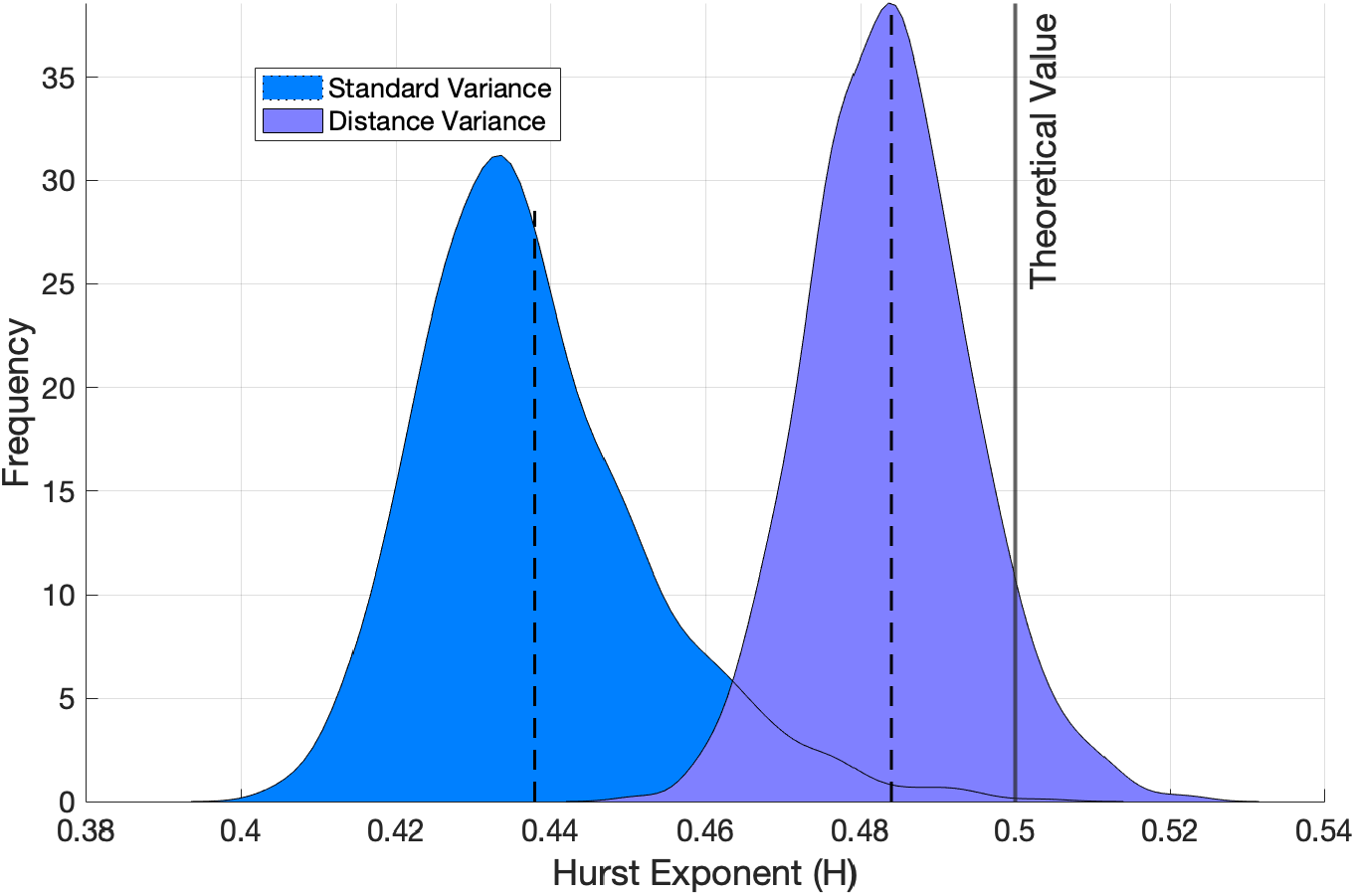
Hurst exponent estimated from the standard and the distance variance-based methods.

A drawback in using the distance variance is its high computational cost [10]. That is, the computational cost for computing the distance variance is *O*(*n*^2^), which limits applicability on large datasets. To lower computational cost, different algorithms have been proposed. For example, the study by Huo and Székely [9] proposed a fast algorithm with a reduced computational complexity of *O*(*n* log *n*). Chaudhuri and Hu [3] also proposed an algorithm with the same computational complexity as that of Huo and Székely. This study, however, uses the algorithm proposed by Chaudhuri and Hu because its implementation is faster.

### Feature Extraction

Instead of using more sophisticated feature selection techniques such as binning and clustering, a simple direct feature extraction is considered in this study. This method uses the whole pupil diameter time series to select discriminatory features with large differences in mean values between cases and controls characterized by Fisher’s criteria (*F*) given in (5).

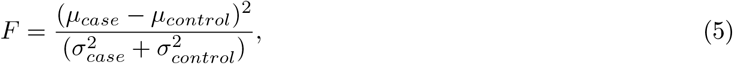

where *μ*_*case*_ and *μ*_*control*_ are the sample means of spectral slopes over the number of subjects in the case and control groups, respectively, and 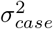 and 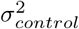 are the corresponding sample variances.

This study, however, uses an evolutionary spectra-based approach for deriving discriminatory features from the raw pupil diameter data. That is, discriminatory features are extracted from a window that moves along the signal with steps of specific size. Spectral slopes in the wavelet spectra calculated for those moving data windows are candidates for discriminatory descriptors. Thus, this approach results in a collection of spectral slopes that are dependent on frequency/scale properties of the signal, but also capture localized features. It is important to note that depending on the step size used to shift the data window along the signal, descriptors could be extracted from overlapping (or non-overlapping) data windows. This study uses non-overlapping data windows. Finally, in order to select the most significant discriminatory descriptors, in this study, we use the Fisher’s criteria (*F*) given in (5).

## Results

This section first presents the data analysis design and then quantifies the self-similar behavior in the pupil-diameter data. Finally, the performance of the proposed method is presented, including comparisons with a feature engineering technique proposed in the literature (readers can find more information about this technique in Das and Khanna [5]).

### Data Analysis Design and Procedure

It is important to note that, in our analysis, we did not include the *ADHD-on-medication* group, as the inclusion of this group introduces bias into the ADHD-diagnosed group [5]. Thus, the dataset used in this analysis is reduced to 50 instances (22 controls and 28 ADHD-diagnosed children off-medication). In addition, we assumed that all participants performed the visuospatial task repeatedly under the same experimental conditions, and thus faced identical difficulty throughout the experiment. The data analysis was performed as Fig. 3 presents, considering one session (20 trials) as a signal of size 165,000. That is, each participant has 8 such signals. MATLAB software was used to perform the data analysis.

**Figure 3:**
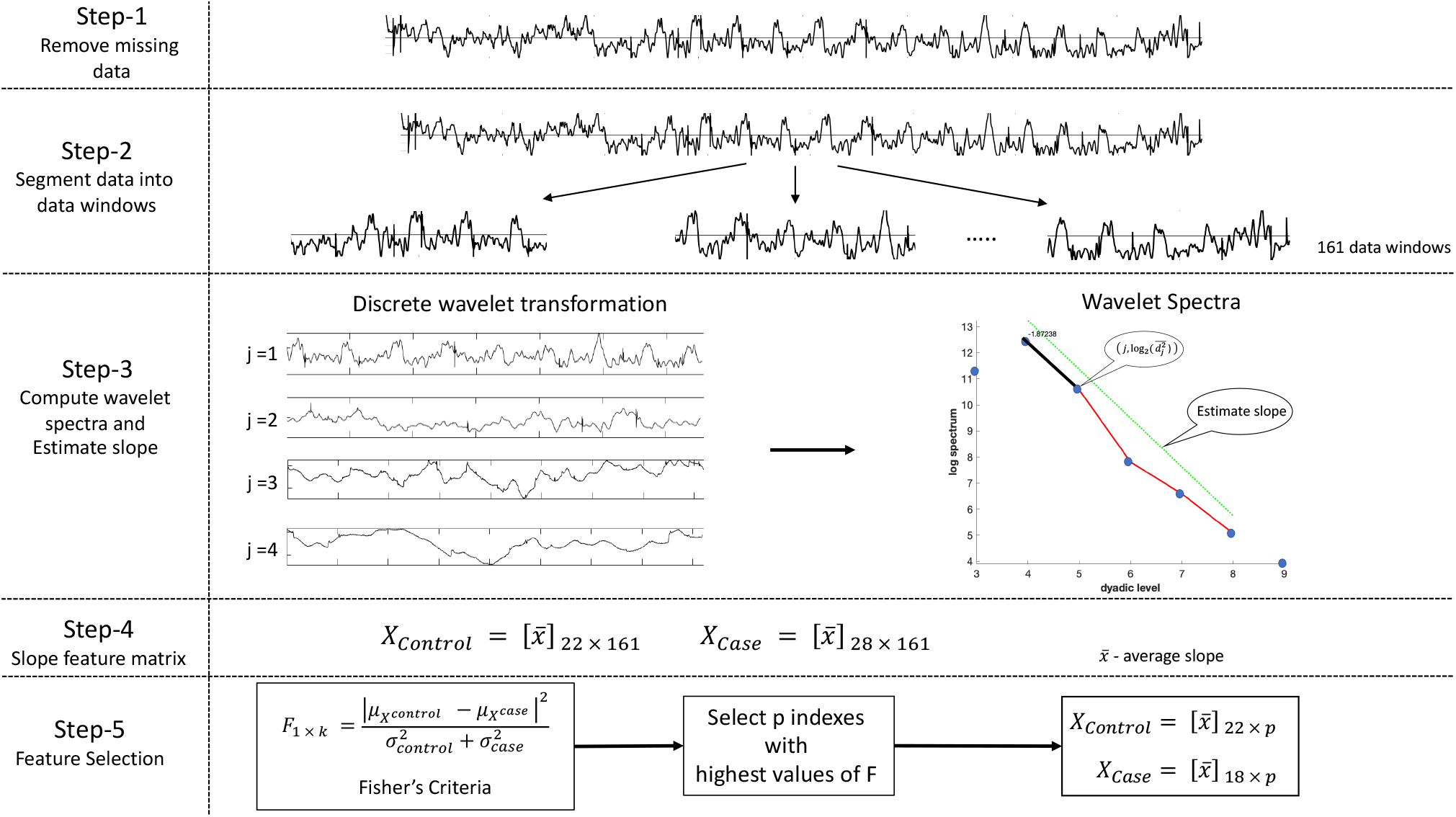
Graphical representation of the data analysis procedure used in this study.

1. **Data Cleaning:** The signals were compacted by removing indices with missing data. The signals with more than 80% missing data were also removed.
2. **Data Segmentation:** Each signal was divided into a set of data windows. For example, if the selected window size was 1024, then 161 data windows per signal were considered, i.e., 165,000/1,024 = 161.
3. **Slope Estimation:** Discrete wavelet transform (DWT) and wavelet spectra were found for each data window in each of 8 signals. We used Daubechies-6 wavelet filter and wavelet decomposition level *J*_0_ = 6. For each participant, this resulted in a matrix spectral slopes of size 8 *× k*, where *k* is the number of data windows. The row average of this slope matrix, 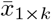), was computed to get the average slope over the eight sessions.
4. **Feature Matrices:** The feature matrices for the controls and ADHD-diagnosed cases (*X*_*Controls*_ and *X*_*Cases*_) were created by using the average spectral slope matrices for individual subjects 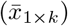 as computed in step (3).
5. **Feature Selection:** The Fisher’s criteria given in (5) was used to select the most significant features from the feature matrices.

The feature matrices formed from the above design were used to feed three classification algorithms: logistic regression (LR), support vector machine (SVM) and K-nearest neighbor (KNN). For model fitting, 67% of the rows were randomly selected from each of the feature matrices for training; the remaining rows were used for testing. In the following sections, classification performance of the three models are discussed by using standard performance metrics sensitivity, specificity, and classification accuracy.

When performing logistic regression (LR), the best threshold on the model predicted probabilities for separation of cases from controls was selected by using the *Youden Index*. This index is defined as the point on the ROC curve (graphical plot of *sensitivity* versus *1-specificity*) which is most distant from the diagonal and computed as 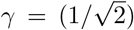 (*sensitivity* + *specificity* − 1) [28]. The threshold that maximizes *γ* is used as the best threshold. Figure 4 shows the ROC curves computed for the feature engineering and self-similarity-based approaches where the best threshold values are 0.22 and 0.33, respectively. The sensitivity, specificity, and accuracy were reported corresponding to these thresholds.

**Figure 4:**
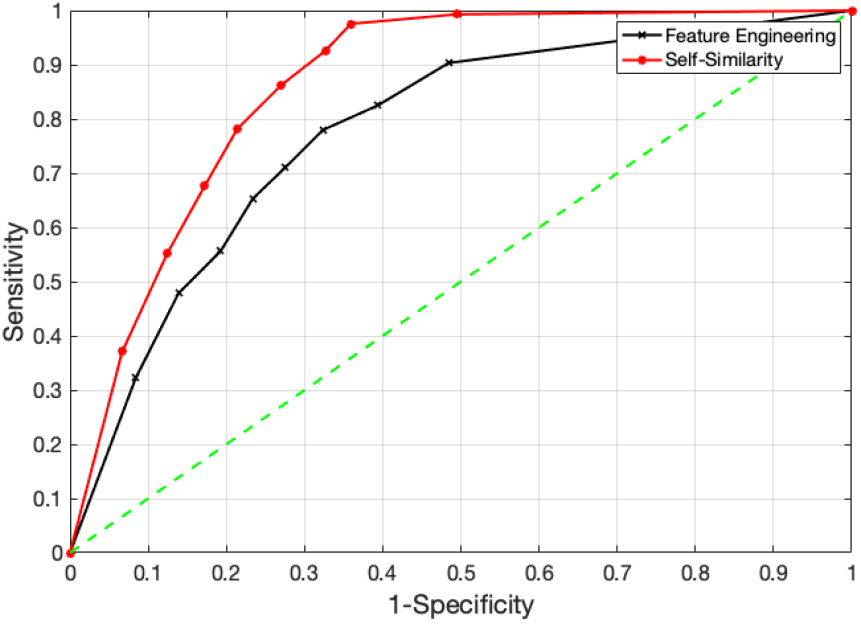
ROC curves obtained by using the feature engineering and self-similarity-based feature matrices.

### Assessment of Self-similar Behavior in Pupil Diameter Data

We investigated the self-similar behavior in the pupil diameter dataset prior to performing our data analysis. For example, Fig. 5a shows a sample signal selected from the controls and cases groups along with their wavelet spectra. These pupil diameter signals posses self-similar properties as their wavelet spectra show a linear decay with scale index. The self-similarity is quantified as the Hurst exponent, which is 0.89 for control and 0.95 for case. When comparing the distribution of the Hurst exponent in the cases and controls, as is seen in Fig. 5b, the pupil diameter of the cases group exhibits relatively less regularity (smaller Hurst exponent), compared to that of the control group. Thus, this reflects that the spectral slope of pupil diameter time series could be utilized as discriminatory descriptors to differentiate ADHD children from controls.

**Figure 5:**
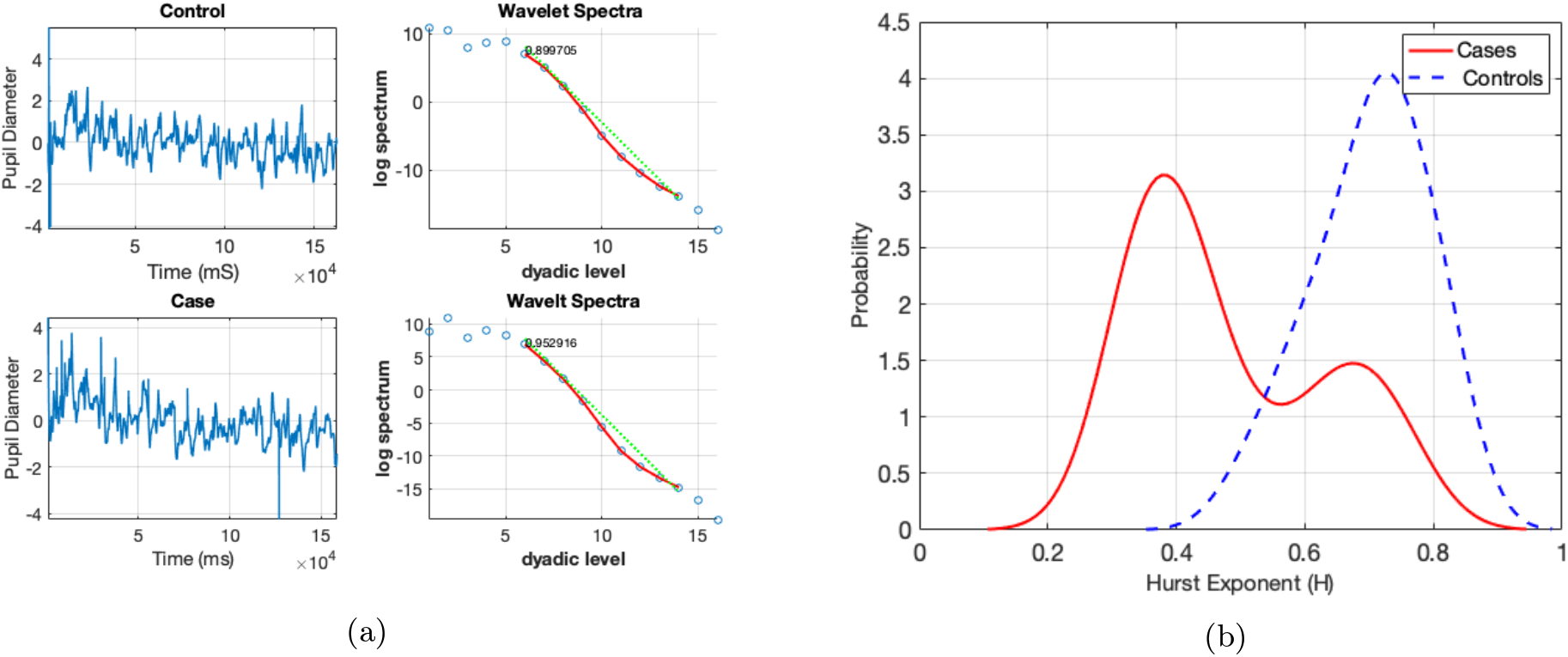
Self-similarity in pupil diameter: (a) a sample pupil diameter signal (one session) and its wavelet spectra along with estimated Hurst exponent and (b) distribution of Hurst exponent for *controls* and *cases* groups.

### ADHD Detection Performance of the Proposed Method

According to the data analysis procedure explained above, classification performance could vary with respect to analysis design settings: data window size and number of classifying features. We first investigated the impact of these settings on classification performance then selected their optimal values to achieve best classification performance. Given that, we selected a wavelet decomposition level of 5. Fig. 6a, 6b, and 6c show variability in accuracy, sensitivity, and specificity with increasing window size and number of classifying features, respectively. Overall, the smaller the window size and the higher the number of classifying features, the better the classification performance. However, an overly small data window size results in a large number of data windows, and hence, data processing could be a relatively expensive computing task. Therefore, data analysis was performed by selecting data windows of size 1024 (log_2_(1024) = 10) and six of the most significant classifying features. Table 1 summarizes classification performance outcomes. The support vector machine model achieved the highest classification accuracy (84.30%) and sensitivity (0.97), followed by the k-nearest neighbor and logistic regression models. The specificity was highest with the logistic regression model (0.76) then the support vector machine and k-nearest neighbor model.

**Figure 6:**
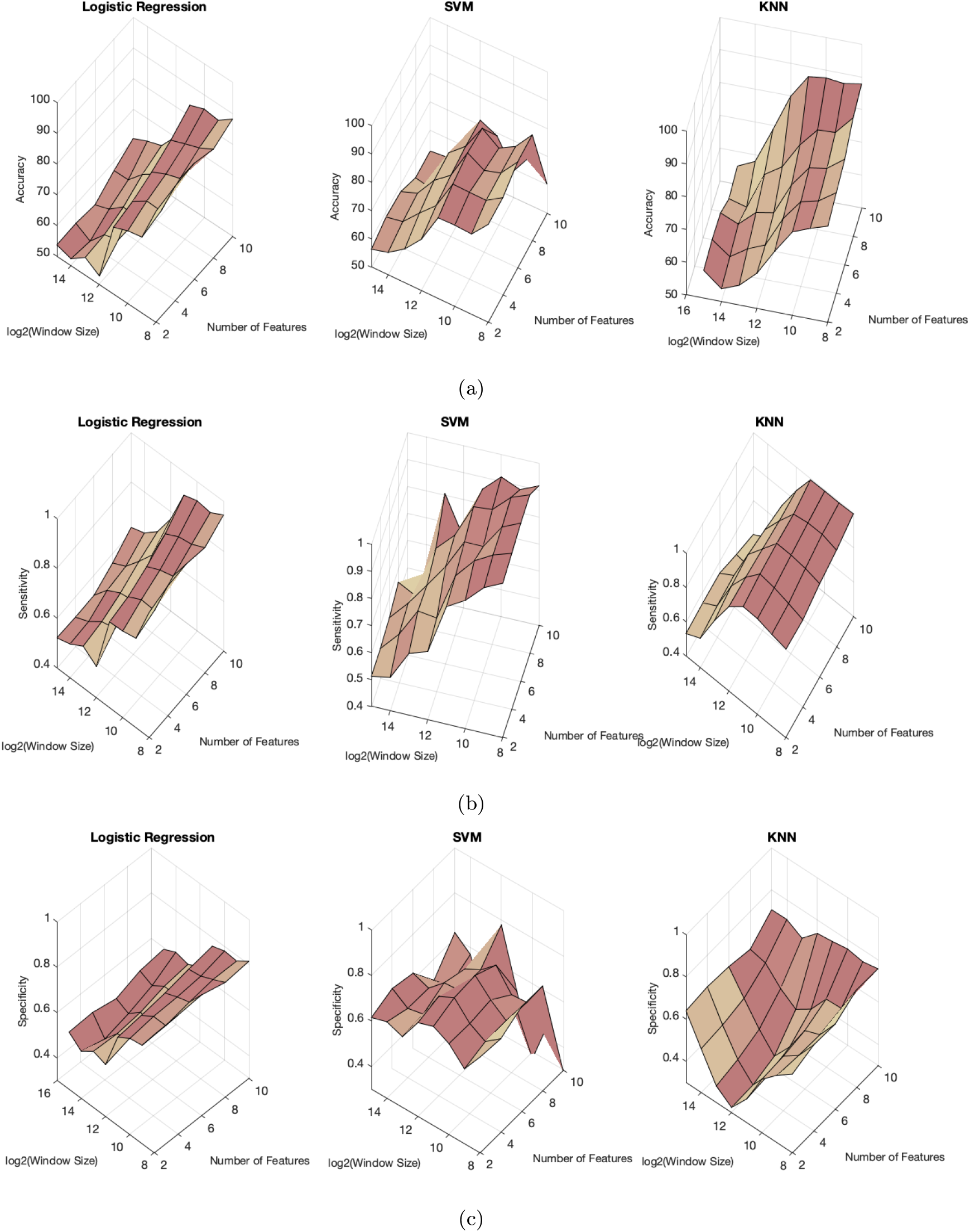
Change in classification performance (a) accuracy, (b) sensitivity and (c) specificity with the window size and the number of features.

**Table 1:**
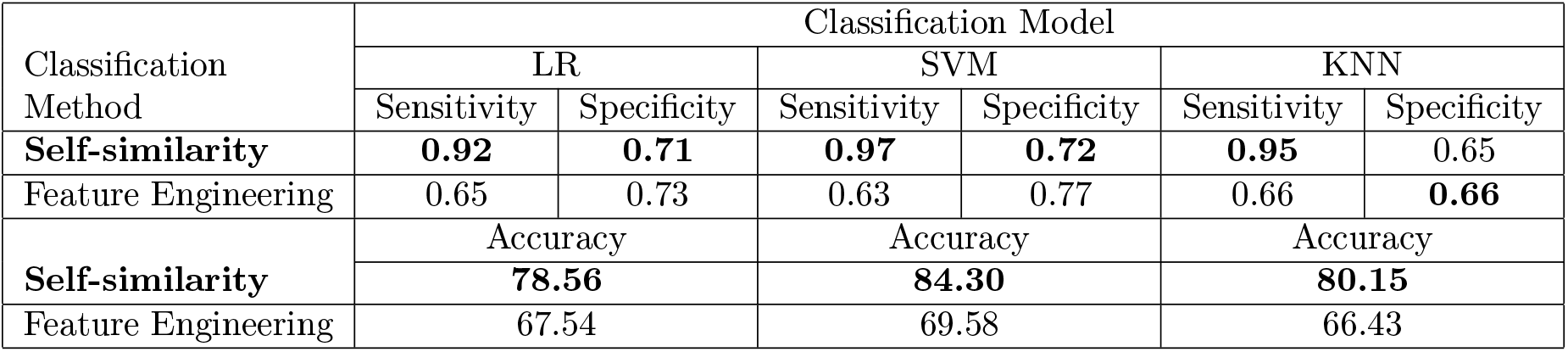
Classification performance of the self-similar behavior-based method (with 1024 window size, and six features and wavelet decomposition levels) and feature engineering method (22 features defined in Das and Khanna [5].

### Performance Comparison with the Existing Methods

We compared performance of the proposed self-similarity-based method with a feature engineering-based method proposed in Das and Khanna [5]. The feature engineering method engineered 22 customized features. Those features were based mostly on standard statistical measures, such as minimum, maximum, mean and median of the z-score values of pupil diameter. The complete list of features used in the feature engineering method can be found in Das and Khanna [5]. The classification performance of these features was explored and compared with the proposed approach (Table 1). Overall, our proposed method shows higher performance compared to the feature engineering method. Thus, the discriminatory descriptors derived though the assessment of self-similar behavior in the pupil diameter data show higher discriminatory power in detecting ADHD compared to the features derived from the original data domain used in Das and Khanna [5].

## Discussion

In this paper, we present a self-similar behavior-based approach to distinguish ADHD-diagnosed children from healthy children using pupil diameter time series. We also present evidence supporting contribution of the proposed approach to improving ADHD detection by comparing its ADHD detection performance with an existing technique.

In the sense of Fig. 5b, pupil diameter time series of the affected individuals were relatively less regular and more erratic compared to non-ADHD children. This is due to the disruption of long-memory correlations in pupillary dynamics affected by external stimuli. This could be attributed to increased inattention in ADHD children as they are typically more prone to distraction by external stimuli compared to non-ADHD children, which would disrupt typical long-memory correlation patterns in pupillary dynamics. Thus, this justifies the potential of the spectral slope as an interpretable discriminatory biomarker to characterize and differentiate subjects with ADHD.

This study shows that self-similar (long range dependence) properties contribute to better extracting of discriminatory information from the pupil diameter time series compared to features derived in the original data domain. That is, for example, the feature engineering method uses standard statistical measures, such as mean, median, and kurtosis, for deriving features. These measures, however, cannot capture hidden patterns that can explain underlying unique behaviors, such as long memory dependence, hidden in high-frequency pupil diameter time series, as is reflected by many studies [6].

A key advantage of the proposed methodology in the present study is that it requires minimal pre-processing. The existing techniques rely heavily on pre-processing pupil diameter data. Since there is no widely accepted standardized methodology for pre-processing, differences in pre-processing could limit the generalizability and reproducibility of informative features. The diagnostic procedure proposed in this study overcomes such limitations as it requires minimal pre-processing. As a result, it can achieve higher classification performance, compared to the ADHD detection techniques that have been published based on the same pupil diameter dataset.

It is already known that the joint use of independently generated features contributes to better explaining natural grouping in data [25]. It is important to note that the features used in the self-similarity-based and feature-engineering method are generated in two independent domains. That is, the self-similarity behavior-based approach uses discriminatory features derived in the wavelet domain, while the feature engineering method relies on features derived in the original data domain. Given the same experimental settings used in Table 1, we investigated the ADHD detection performance of different combinations of features selected from these two methods with respect to accuracy. However, we could not observe an improvement with all three classification models. The LR model achieved approximately 5% improvement in accuracy, but SVM and KNN models declined accuracy for nearly 6%. Further analyses are essential to reveal reasons for this decline in classification accuracy.

It is important to note that, based on our pupil diameter data analysis, the most informative information that could be used to detect ADHD condition is concentrated towards the end of the signals. That is, as can be seen in Figure 7, the most discriminatory features selected through the proposed approach are from the latter part of the signal. This could be due to the decrease in attention and increase in hyperactivity and impulsivity that is typical of ADHD-diagnosed participants over the series of experimental trials. However, this preliminary finding warrants more investigation.

**Figure 7:**
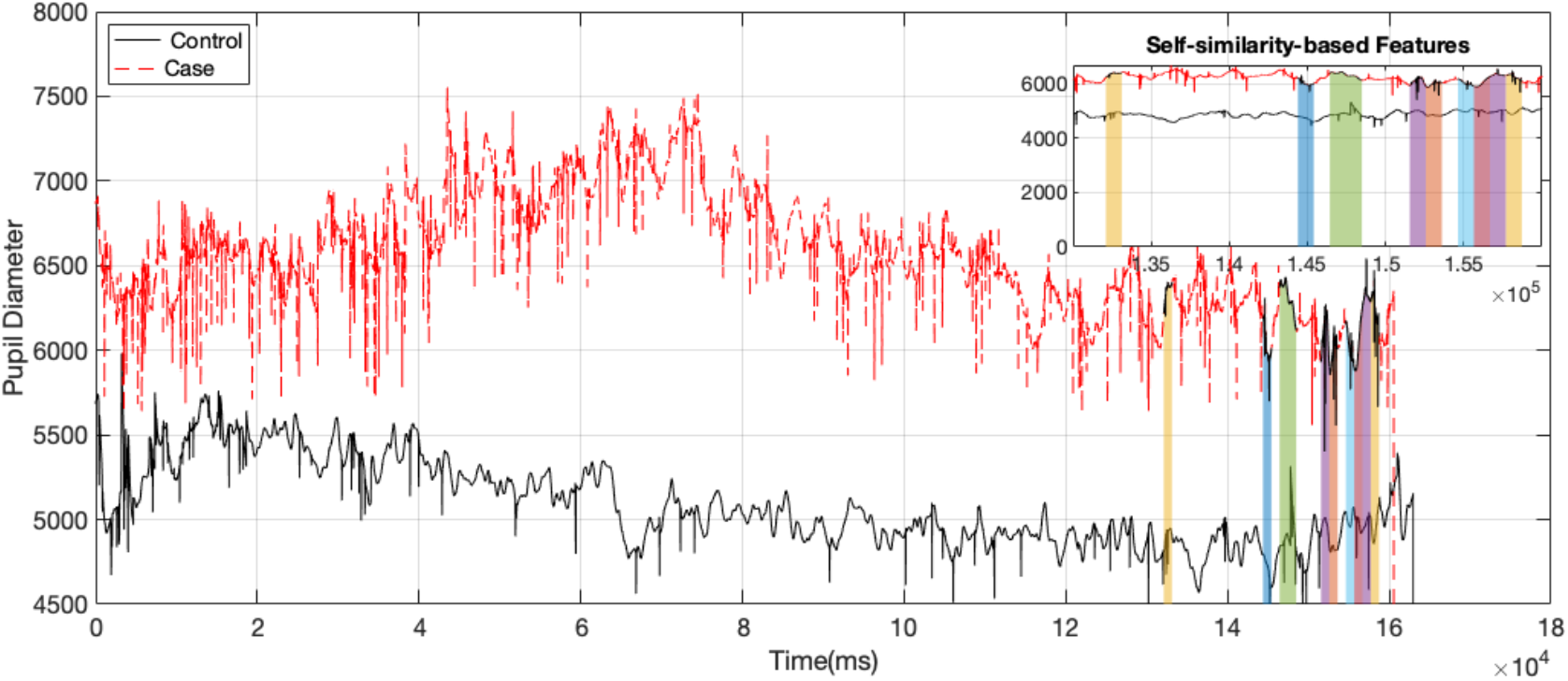
An example pupil diameter signal selected from the normal and ADHD-diagnosed (without medication) group, and the most discriminatory features selected by the self-similarity-based approach.

## Conclusions

This study proposed a method to detect ADHD based on the self-similar property of pupil diameter time series. Self-similarity was assessed in the wavelet domain by computing wavelet spectra of pupil diameter time series. The spectral slope of the wavelet spectra was used to derive discriminatory features via a rolling window-based approach. The performance of the derived features was explored then compared with an existing feature engineering method. Application of this method on pupil diameter data showed that the proposed method achieved greater than 78% accuracy in detecting ADHD, which is higher than that of the existing method, which involves feature engineering in the original data domain. The discriminatory features derived via the wavelet-based self-similarity analysis improves ADHD detection performance. Therefore, the self-similar property of pupillary dynamics could be used as a potential biomarker for automated detection of ADHD effectively.

